# A Microfluidic Blood Vessel-On-Chip Model of Thrombosis

**DOI:** 10.1101/2025.07.10.663679

**Authors:** Josefin Jansson-Edqvist, Tatiana Mencarini, Claire Peghaire, Matthew Dibble, Josefin Ahnstroem, Charis Pericleous, Alberto Redaelli, Joseph van Batenburg-Sherwood, Anna M. Randi

**Author notes:** Corresponding Authors: Anna M. Randi MD PhD FAHA, Imperial Centre for Translational and Experimental Medicine, Imperial College London, NHLI Vascular Sciences, Hammersmith Hospital, Du Cane Road London W12 0NN, Joseph van Batenburg-Sherwood, Department of Bioengineering, Imperial College London, Sir Michael Uren Building 86 Wood Lane, London W12 0BZ. University of Bordeaux, France (Claire Peghaire).

## Abstract

**Background:** Thrombus formation is regulated by the interplay between endothelial cells (EC), platelets and coagulation factors. However, most *in vitro* assays used to study thrombosis and develop anti-thrombotic therapies do not include the endothelium.

**Objective:** To develop a thrombus-on-chip model that includes endothelium and whole blood and allows for manipulation of extracellular matrix (ECM), shear stress and the addition of multiple cell types.

**Methods:** A cylindrical vessel was created in a collagen matrix using the needle-based fabrication technique in a microfluidic device. Human umbilical vein EC (HUVEC) or endothelial colony-forming cells (ECFC) were cultured in the channel, with continuous monodirectional media turnover. Immunostaining and permeability measurement validated confluence and integrity of the monolayer. To investigate thrombosis, whole blood from healthy donors was perfused through TNF-α-activated or untreated EC-lined vessels. Image analysis and time-lapse microscopy were used to quantify labelled platelet adhesion and fibrin deposition.

**Results:** TNF-α treatment resulted in increased platelet adhesion and fibrin deposition compared to control vessels. TNF-induced endothelial activation was confirmed by upregulation of adhesion molecules ICAM-1, E-selectin and tissue factor (TF). Thrombus formation in TNF-α treated vessels was inhibited by an anti-TF antibody. In ECFC vessels, platelet adhesion and fibrin deposition were comparable to HUVEC, supporting feasibility of patient-based studies.

**Conclusions:** We developed a perfused thrombus-on-chip model that combines key elements of thrombus formation including endothelium. The model is amenable to independent control of microenvironmental stimuli, crosstalk with tissue-specific cells, and the inclusion of patients’ own cells and blood for precision medicine studies.

**ESSENTIALS:**

- The endothelium is a key contributor to thrombosis
- Standard in vitro methods to study thrombosis do not include the endothelium
- We have developed a thrombus-on-a-chip model of thromboinflammation to measure live platelet adhesion and fibrin deposition on activated endothelium
- Using endothelial colony forming cells (ECFC), we show that the method is suitable for patient studies with autologous endothelium and blood

## INTRODUCTION

Haemostasis is the physiological response to vessel wall injury, a protective mechanism to prevent excessive blood loss. However, pathological clot formation can result in thrombosis, a leading cause of morbidity and mortality across the world. The search for better diagnosis and treatments for thrombotic diseases remains a priority.

Physiologically, haemostasis is triggered by vessel wall injury, which exposes subendothelial collagen matrix and tissue factor (TF) and initiates clot formation [1], [2], [3]. Simultaneously, activation and adhesion of platelets to collagen fibrils and/or von Willebrand factor (VWF) released from activated EC results in the formation of the platelet plug, followed by platelet degranulation and further platelet aggregation. Secondary haemostasis involves the coagulation cascade and activation of plasma clotting factors, leading to fibrin formation stabilising the clot [4].

Multiple factors can disrupt the complex balance of haemostasis and lead to thrombosis. Abnormalities or an increase in circulating coagulation and fibrinolytic factors, which increase the risk of thrombotic events, are well described [5]. However, despite the equal systemic exposure of the vasculature to circulating factors, thrombotic diseases tend to prefer specific sites. The reasons for this are unclear but suggest local factors, likely related to the activation state of the endothelium. In support of this concept, endothelial activation caused by inflammation and/or disturbance of blood flow has been widely correlated with the pathogenesis of cardiovascular diseases, including arterial and venous thrombosis [6], [7], [8].

In homeostatic conditions, the endothelium presents as an anti-coagulant surface; this switches to a pro-coagulant surface when required and then reverts to its homeostatic state. Pathological activation of the endothelium results in loss of its anti-thrombotic properties. Endothelial function and phenotype are influenced by multiple factors in the microenvironment, including the extracellular matrix (ECM) [9], substrate stiffness [10], [11], [12], blood flow [6], [7], [8] and the crosstalk between the EC and other vascular cell types, such as fibroblasts, smooth muscle cells and pericytes [13], [14]. Little is known about the contribution of these factors to thrombosis. These pathways may be of crucial importance to understand local mechanisms of thrombosis, and thus offer new targets for diagnosis, biomarkers and anti-thrombotic therapies.

Most diagnostics and therapies focus on circulating factors rather than aiming at reducing the pro-coagulant activity of the endothelium. This may be partly due to the lack of *in vitro* models of thrombosis that incorporate vascular cells. Encompassing all the key factors contributing to thrombus formation in a single *in vitro* model is challenging. Standard 2D cell culture techniques cannot reproduce the complexity of the physiological microenvironment; hence new models that capture multiple features to bridge the gap between *in vivo* and *in vitro* models are required. Lately, a significant effort has focused on the development of cell-based *in vitro* microfluidic or ‘organ-on-chip’ models that include ECs, ECM, other vascular cells and/or whole blood and shear stress [15], [16], [17]. As recently reviewed [18], an ideal ‘organ-on-chip’ vascular model would allow for the incorporation of EC, mural and other vascular cells, an elastic interface that can allow for ECM-EC signalling, and a set up that allows the generation of physiological and pathophysiological flow. Furthermore, investigation of thrombus formation also requires the ability to perfuse blood and record thrombus formation by time-lapse microscopy. The geometry of these devices is crucial, since red blood cells promote the margination of platelets, increasing the frequency of interactions with EC [19]. Direct contact between EC and ECM creates an environment close to physiological stiffness, also capable of accommodating other cells lineages and blood perfusion. Recently, several models have been developed to investigate thrombus formation *in vitro* [15], [20], [21], [22], [23], [24], [25]; however, the challenge remains to develop a model with relevant geometry that can incorporate all variables.

The ambitious goal of precision medicine requires access to patients’ own cells and biological samples. For the vascular endothelium, this is now possible thanks to endothelial colony-forming cells (ECFC), circulating progenitors isolated from donors’ peripheral blood that can be expanded in culture for multiple applications [26]. A recent study showed dysregulation of the TNF pathway in ECFC from patients with unprovoked thromboembolism [27], supporting the value of this approach in investigating hidden causes of thrombosis.

In this study, we develop a vessel-on-chip model of thrombosis that incorporates the above-mentioned key components of haemostasis and allows for precision medicine approaches. A needle-based fabrication technique was used to create a 3D cylindrical channel embedded in a collagen matrix, which was then lined with EC. The model allows for direct contact between EC and ECM, exposing cells to physiological stiffness; it includes perfusion and exposure of EC to shear stress; it allows live imaging of thrombus formation, inclusion of activators and inhibitors, and molecular analyses of ECs, including patients’ own EC, for the development of a personalised model of thrombosis. The model is amenable for future development to include vascular mural and tissue cells, different ECM and different shear stress.

## METHODS

### Vessel-on-chip fabrication

Model fabrication was based on needle casting [28] to generate a cylindrical perfusable microfluidic channel and involved three steps: i) polydimethylsiloxane (PDMS) is cast in a 3D printed mould to create the chip, ii) the chamber in the chip is filled with collagen around a needle, which is withdrawn to create the lumen, iii) the lumen is endothelialised and cultured to achieve a mature monolayer.

#### Manufacturing the chip

A mould (Figure 1A) was printed with a polyjet 3D printer (Objet 30 Prime, Stratasys, USA). The outer rim of the mould was designed with two ECM ports and two vessel ports, all leading to a chamber for the ECM (diameter=5 mm, height=1.3 mm). Polytetrafluoroethylene (PTFE) tubing (OD=0.6 mm, Sigma Aldrich, USA) was then inserted into the ECM ports through two holes in the chamber region. To create the main vessel structure, a 300 µm diameter stainless-steel acupuncture needle (Seirin, Japan) was inserted through the ECM chamber, through tubing at the vessel inlets, leaving approximately 2 mm between the tubing and the ECM chamber (Figure 1A). PDMS (Sylgard 184, Dow Corning, USA) was mixed at a 10:1 elastomer to curing agent ratio, poured into the mould and polymerised at 65°C for two hours. After curing, the tubing and needle were removed, leaving hollow inlets in the PDMS and a cavity for the ECM. The PDMS support structure was then removed from the mould and sealed with a glass coverslip, which was bonded to the PDMS using oxygen plasma treatment of both the glass and PDMS surfaces, followed by post-treatment baking at 65°C for one hour (Figure 1B).

**Figure 1:**
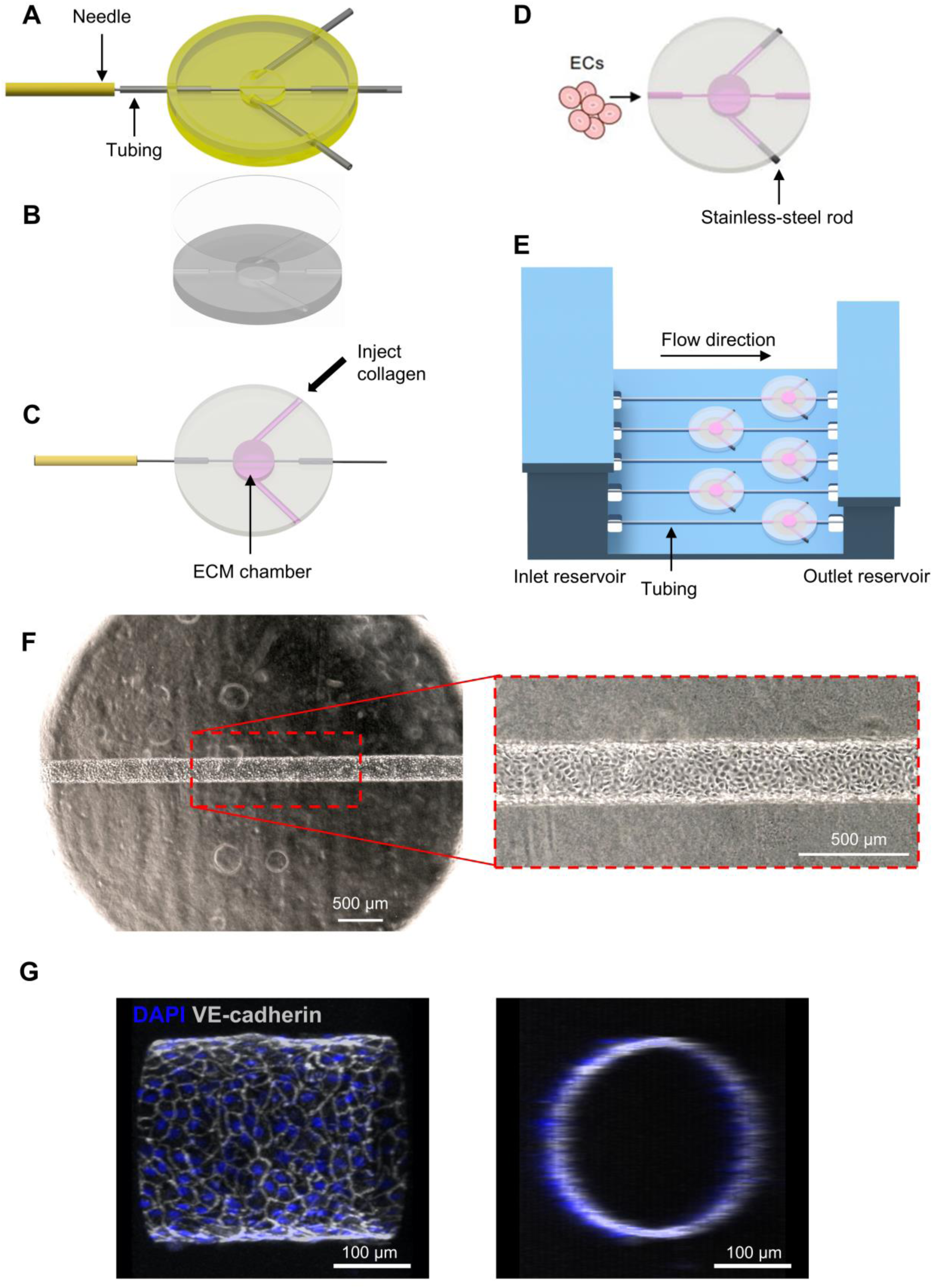
Constructing the model. A-D: Fabrication process of the microfluidic chip. A: Mould preparation, tubing and needle positioning. B: Cured PDMS extracted from the mould and circular glass coverslip. C: Sterilised needle inserted in the central channel and collagen pipetted from the lateral port. D: After collagen reticulation and needle extraction, a circular lumen is formed in the gel. Stainless-steel rods are positioned to block the lateral ports. Endothelial cells are seeded in the empty circular channel. E: Schematic of the setup: 3D-printed unit housing five microfluidic devices, connected to tubing for continuous media turnover. F: Phase contrast image showing a confluent microvessel lined with HUVECs 4-5 days post seeding, featuring a typical cobblestone morphology. Inset shows enlarged image the central portion. G: Confocal images of the vessel immunostained against DAPI (blue) and VE-cadherin (white) 5 days post seeding. The cross section confirms the complete coverage and circularity of the microvessel lumen.

#### Creating the lumen of the vessel-on-chip

All inlets and the ECM chamber were sterilised with 70% ethanol and washed with PBS. To promote adhesion of the collagen gel to PDMS, the ECM chamber and ECM inlets were coated with neutralised collagen solution (rat-tail collagen I 50 µg/ml, Corning, USA) overnight at room temperature. After removing the collagen solution, the ECM chamber and all inlets were filled with PBS, and an identical 300 µm stainless-steel acupuncture needle, sterilised with 70% ethanol, was inserted through the vessel inlet into the ECM chamber. The PBS was then aspirated from the ECM inlets.

Collagen was neutralised in HEPES (1M, Sigma Aldrich, USA) and NaHCO_3_ (37 g/l reconstituted in MilliQ water, Sigma Aldrich, USA) at an 8:1:1 ratio on ice to slow down polymerisation during preparation. The neutralised collagen gel (4 mg/ml) was injected through one ECM inlet, filling the ECM chamber around the needle (Figure 1C) and left to polymerise at 37°C overnight. Following collagen polymerisation, the needle was withdrawn, leaving a straight hollow cylindrical channel, surrounded by collagen in the ECM chamber. Stainless-steel rods, sterilized with 70% ethanol, were inserted to seal the ECM inlets to inhibit flow (diameter=0.8 mm, length: ∼2-3 mm cut from a wire, RS Components Ltd., UK ) (Figure 1D).

#### Endothelialising the lumen

Two types of EC were used in the study. Human umbilical vein ECs (HUVEC, from Lonza, UK: pooled, 10 donors) were used at passage 4-6. Alternatively, endothelial colony forming cells (ECFC) from healthy donors were isolated from blood samples as described previously [29]. ECFC from five individual donors were combined at passage 4 to generate a pool, which was further expanded for one passage to passage 5. Written informed consent for ECFC isolation and use was obtained under the Imperial College Healthcare Tissue Bank (HTA licence 12275; REC approval 17/WA/0161).

HUVEC or ECFC in EC growth medium-2 (EGM2, Lonza, UK) were seeded in the lumen at a density of 5×10^6^ cells/ml, by injecting 2.5 µl of cell suspension through the vessel inlet (Figure 1D). The device was incubated at 37°C for 30 minutes to allow cell adhesion to the collagen; then the device was rotated 180° and the cell seeding process was repeated to ensure cell coverage.

To maintain cell viability and allow for parallelisation, the device was connected via PTFE tubing (ID=0.3 mm, OD=0.76 mm, Adtech Polymer Engineering Ltd, UK) to a perfusion unit, fabricated from nylon (PA2200) using a SLS 3D printer (Forge Labs, Canada) (Figure 1E). To prevent leaking, the 3D printed reservoirs were coated with PDMS and incubated overnight in an oven at 65°C for polymerisation. The perfusion unit has the dimensions of a rectangular tissue culture plate, in order to fit on a microscope stage and allows optical access through holes in the base. The unit has five separate inlet and outlet reservoirs to enable culturing of five vessels in parallel. Hydrostatic pressure-driven flow from the inlet reservoirs to the outlet reservoirs was used to ensure a continuous renewal of culture media for the cells. As fluid flows from the inlet to the outlet reservoir, the height of fluid in the inlet reservoir decreases and the height of fluid in the outlet reservoir increases; this reduces the pressure gradient driving the flow and limits the duration that a given flow rate can be maintained. To allay this effect, the outlet reservoirs were designed with an overflowing feature to maintain fluid height (see Appendix A1). This enabled continuous media turnover of at least one vessel volume per minute, with media replacement required only once per day. The shear stress induced by continuous flow through the vessel was low throughout the culture period (<0.3 dyne/cm^2^) (Appendix A1). The perfusion unit was sterilized by autoclaving before use. Cell culture media was replaced once per day with 3 ml of EGM2 supplemented with 25 mM HEPES, pre-conditioned overnight to incubator conditions to minimise air bubble formation in the tubing and vessel. Endothelialised vessels were cultured for up to 9 days prior to IF or thrombosis assays.

### Immunofluorescence (IF) staining

Fixation, permeabilisation, blocking and PBS washes were performed by perfusion; all steps in the protocol were performed on the same day and at room temperature. Vessels were fixed with 4% paraformaldehyde solution for 15 minutes and permeabilised with 0.2% Tween-20 for 20 minutes, followed by 30 minutes blocking with 10% bovine serum albumin (BSA). The primary and secondary antibodies were injected into the vessel through the vessel inlet. The primary antibody for VE-cadherin was delivered in two steps, injecting 5 µl antibody solution (diluted in 1% BSA) and incubating for 45 minutes and then rotating the device 180°, adding an additional 5 µl antibody solution and incubating for another 45 minutes. The vessel was then washed with PBS for 10 minutes followed by incubation with the secondary antibody and DAPI (nuclear stain), which were delivered using the same method as for the primary antibody but with 30-minute incubations. The vessel was washed again with PBS for 10 minutes prior to microscopy or storage at 4°C.

### Thrombosis assay

Blood was collected ∼30 minutes prior to starting the thrombosis assay from healthy volunteers (male and female, age 25-40) who had not taken any pain relief medication known to affect coagulation for the past two weeks. Blood was collected in 3.2% trisodium citrate in a syringe with a 21G butterfly needle. The first 3 ml of blood were discarded and the sample was mixed by slowly inverting the collection tube to minimise shear-induced platelet activation. Blood was stored at room temperature and used within 3h. Platelets were labelled using a phycoerythrin-conjugated anti-human CD41 antibody (BioLegend, USA) at a ratio of 1:100 antibody:blood. Alexa Fluor 647-conjugated fibrinogen from human plasma (Thermo Fisher Scientific, USA) was added to blood at a ratio of 1:200. Five minutes prior to perfusion, blood was recalcified by adding a recalcification buffer (100 mM CaCl_2_ and 75 mM MgCl_2_ in PBS) at a ratio of 1:10 [30].

The perfusion unit was placed on a confocal microscope (LSM 510, Zeiss, Germany) for real-time monitoring. One vessel at a time was perfused with blood using a 1 ml plastic syringe and a syringe pump (Model X, Harvard Apparatus, USA); with the PTFE tubing in the perfusion unit connected to the syringe via a custom PDMS connector. Recalcified blood was perfused at 40 µl/min for 20 minutes, inducing a shear stress of ∼10 dyne/cm^2^ Every 2 minutes, a z-stack of 14 images was captured (each image was 512×1024 12-bit pixels) at the centre of the bottom surface of the vessel using a 10x objective, keeping gain, laser power and pinhole settings constant for all experiments.

Platelet adhesion and fibrin deposition were quantified by analysing the total number of pixels in the z-stack with positive fluorescent intensity and normalising to the captured vessel surface area to account for variability between vessels (Appendix A3). In selected experiments, an anti-Tissue Factor (TF) antibody (mouse monoclonal, IgG_2a_, Enzyme Research Laboratories) or IgG control were used at 25 µg/ml, diluted in cell culture media to the desired concentration and added to the vessel for 20 minutes after TNF-treatment and prior to blood perfusion.

### RNA extraction and RT-qPCR

HUVEC were lysed within the microfluidic device by pipetting Arcturus PicoPure Extraction buffer (Applied Biosystem, Thermofisher, USA) into the vessel and RNA was extracted using the Arcturus PicoPure RNA isolation kit. For ECFC experiments, the collagen gel was removed from the ECM chamber by breaking the coverslip and digested at 37°C in 1 ml of collagenase A solution (Sigma, 2 mg/ml) reconstituted in EGM2 cell culture media for 2-5 minutes until it was no longer visible. The sample was centrifuged at 1200 rpm for 5 minutes at room temperature, after which the collagenase solution was discarded, cells were lysed with Arcturus PicoPure Extraction buffer, and RNA extracted using the Arcturus PicoPure RNA isolation kit.

Total RNA was reverse transcribed into cDNA using Superscript III Reverse Transcriptase (Invitrogen, USA). ICAM1, thrombomodulin (TM), tissue factor (TF), von Willebrand factor (VWF) and E-selectin were measured in HUVEC (n=8 vessels); ICAM1, TM, TF and VWF were also measured in ECFC (n=3 vessels). Quantitative real-time PCR was performed using PerfeCTa SYBR Green Fastmix (Quanta Biosciences, USA) on a CFX96 system (Bio-Rad, UK). Gene expression values were normalised against the housekeeping gene, glyceraldehyde 3-phosphate dehydrogenase (GAPDH). Fold differences were calculated using the ΔΔCt (delta-delta cycle threshold) method, averaging across all controls.

### Permeability

Permeability was measured by replacing the media in the inlet reservoir with 3 ml TRITC-dextran solution (65-85 kDa, tetramethylrhodamine isothiocyanate dextran, Sigma Aldrich, USA), reconstituted in the HEPES-supplemented EGM2 cell culture media to a final concentration of 2 mg/ml). The 3D-printed perfusion unit was placed on the confocal microscope stage and imaged by time-lapse microscopy (7 seconds between images, 65 images, size: 1024×1024 pixels). Microscopy settings were kept the same for all experiments. Following confocal imaging, the dextran solution in the inlet reservoir was replaced with HEPES-supplemented EGM2 cell culture media (pre-conditioned in the incubator) and the perfusion unit was placed back in the incubator for continued cell culture. Permeability assays were performed on different days, to capture different stages of cell confluency and maturation, and to evaluate the progression of monolayer formation and the integrity of the vessel. 14 vessels in total were measured, with 6 having longitudinal measurements to evaluate progression.

Permeability (P_s_) for a vessel at a given timepoint was estimated by quantifying the rate of change of fluorescent intensity inside the ECM over time, using methods similar to those previously reported with minor modifications to the implementation (see Appendix A2 for details). Permeability values are reported in µm/s (equivalent to 10^−4^ cm/s).

### Co-culture with perivascular cells

To investigate the feasibility of co-culture in the vessel-on-chip model, red fluorescent protein (RFP)-labelled human brain microvascular pericytes (HBMVP) (Angio-Proteomie, MA, USA) were seeded in the chip together with HUVEC. HUVEC were seeded first on one side and allowed to adhere for 30 min at 37°C, then the devices were rotated 180° and a HBMVP/HUVEC mix (1:5 ratio) was seeded, resulting in a total cell ratio of 1:10. The chips were incubated upside-down at 37°C for 4h before connection to the culture unit. Confluency was verified by phase contrast microscopy, then the vessels were fixed and IF staining for DAPI and VE-cadherin was performed as described above (see “Immunofluorescence (IF) staining”).

### Statistical Analysis

Appendix A4 contains full details of the statistical analysis. Briefly, MATLAB was used for all statistical analyses. Pairwise comparisons were carried out using two-sample t-tests, with Bonferonni-Holm approach to account for multiple comparisons and reduce the likelihood of a false positive. All p-values are reported as <0.05, <0.01 or <0.001.

## RESULTS

### Characterisation of the collagen-embedded vessel monolayer

Monitoring the microvessels by phase contrast microscopy over time revealed that EC reached confluency with the typical monolayer cobblestone morphology 4-5 days post seeding (Figure 1F). IF staining for the junctional molecule VE-cadherin (day 4-5) confirmed the presence of a continuous confluent monolayer lining the cylindrical space, an open lumen and adherens junctions with no visible gaps (Figure 1G). We therefore proceeded with the functional assay.

### Thrombosis assay

HUVEC- and ECFC-lined vessels were perfused with medium containing TNF-α (10 ng/ml) or control for 4 hours. Freshly collected whole blood, labelled with fluorescently conjugated anti-human CD41 antibody and with fluorescently conjugated fibrinogen, was perfused into the vessels; platelet capture and fibrin formation were studied via real-time confocal imaging with z-stacks captured every 2 minutes over a 20 minutes interval (see Methods for details).

#### HUVEC-lined vessels

Representative images of thrombus formation at the end point of the assay (20 minutes of perfusion) showed greater platelet adhesion and fibrin deposition in TNF-α stimulated vessel (Figure 2A, bottom panels) compared to untreated vessel (Figure 2A, top panels). Some platelet adhesion was observed in untreated vessels, possibly due to low-level platelet activation induced by blood sampling; minimal levels of fibrin were detected in untreated samples. Quantification shows a significant increase in platelet adhesion (Figure 3A) and fibrin deposition (Figure 3B) in TNF-α-treated vessels compared to untreated vessels (p<0.01 for platelet adhesion and p<0.001 for fibrin deposition). Variability in the signal prompted us to investigate reproducibility within the same blood donor, by testing the same blood sample on different vessels (Figure 3A-B, inset). These showed some variability in the platelet adhesion, and more so in the fibrin accumulation data, suggesting that this parameter might be more sensitive to variability between blood vessels.

**Figure 2:**
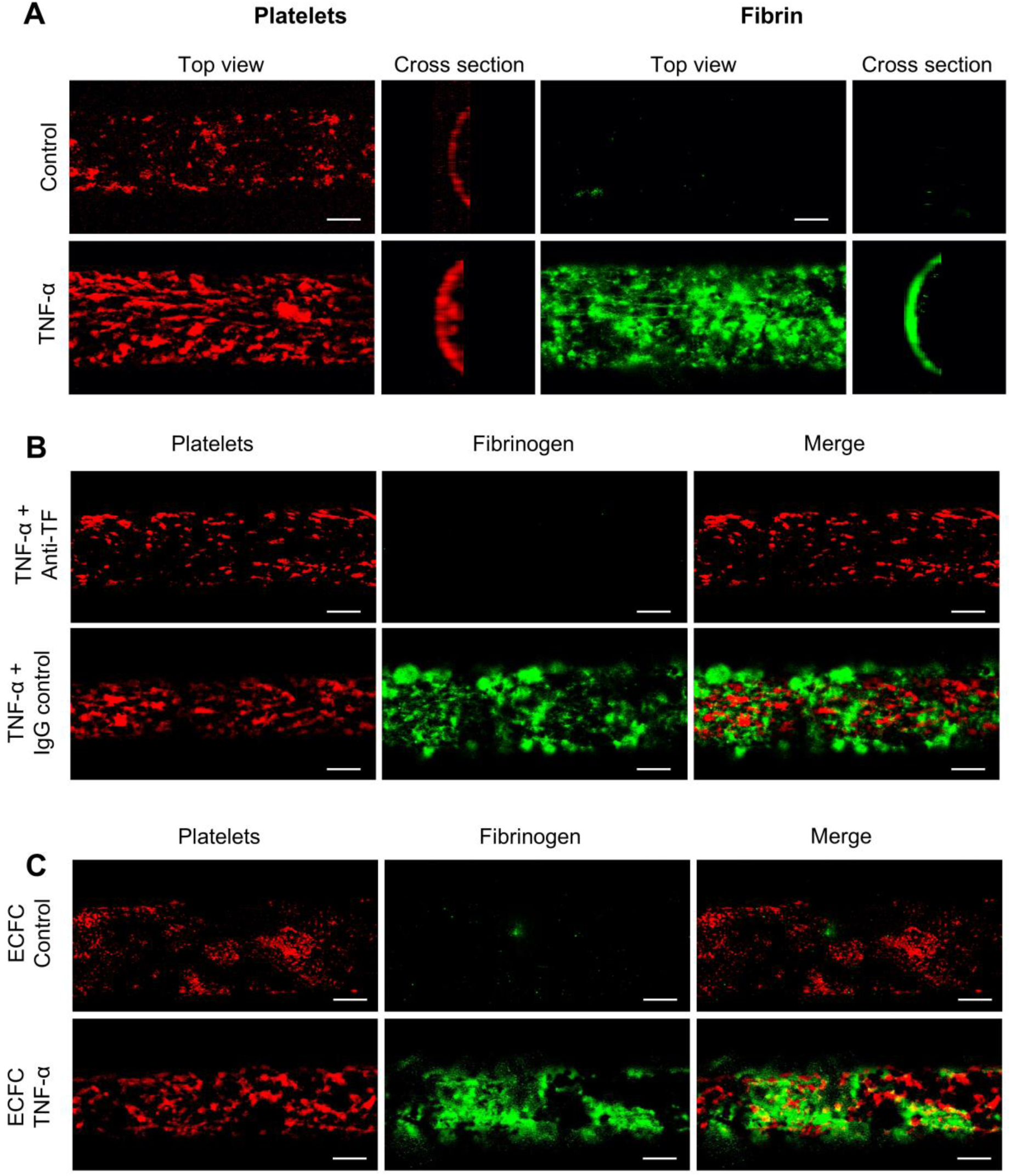
Thrombosis assay: platelet adhesion and fibrin deposition. Representative data (z-stack projections) of platelet adhesion and fibrin deposition from the 3D thrombosis assay after 20 minutes of perfusion. Scale bars = 100 µm. A: HUVEC-lined vessels (control [top] and TNF-α stimulated vessels [bottom]). The TNF-α stimulated vessels show increased platelet adhesion and fibrin deposition compared to a background platelet signal and almost no fibrin deposition in the controls. B: HUVEC-lined vessels stimulated with TNF-α in the presence of anti-TF antibody or IgG control. Blocking TF completely inhibits fibrin deposition induced by TNF, whilst IgG control has no effect. The Anti-TF Ab also decreases TNF-α induced platelet deposition. C: ECFC-lined vessels in control and TNF-α stimulated conditions. Similar to HUVEC (A), TNF-α activation of ECFC results in significant platelet adhesion and fibrin deposition compared to a background platelet signal and almost no fibrin deposition in the untreated vessels.

**Figure 3:**
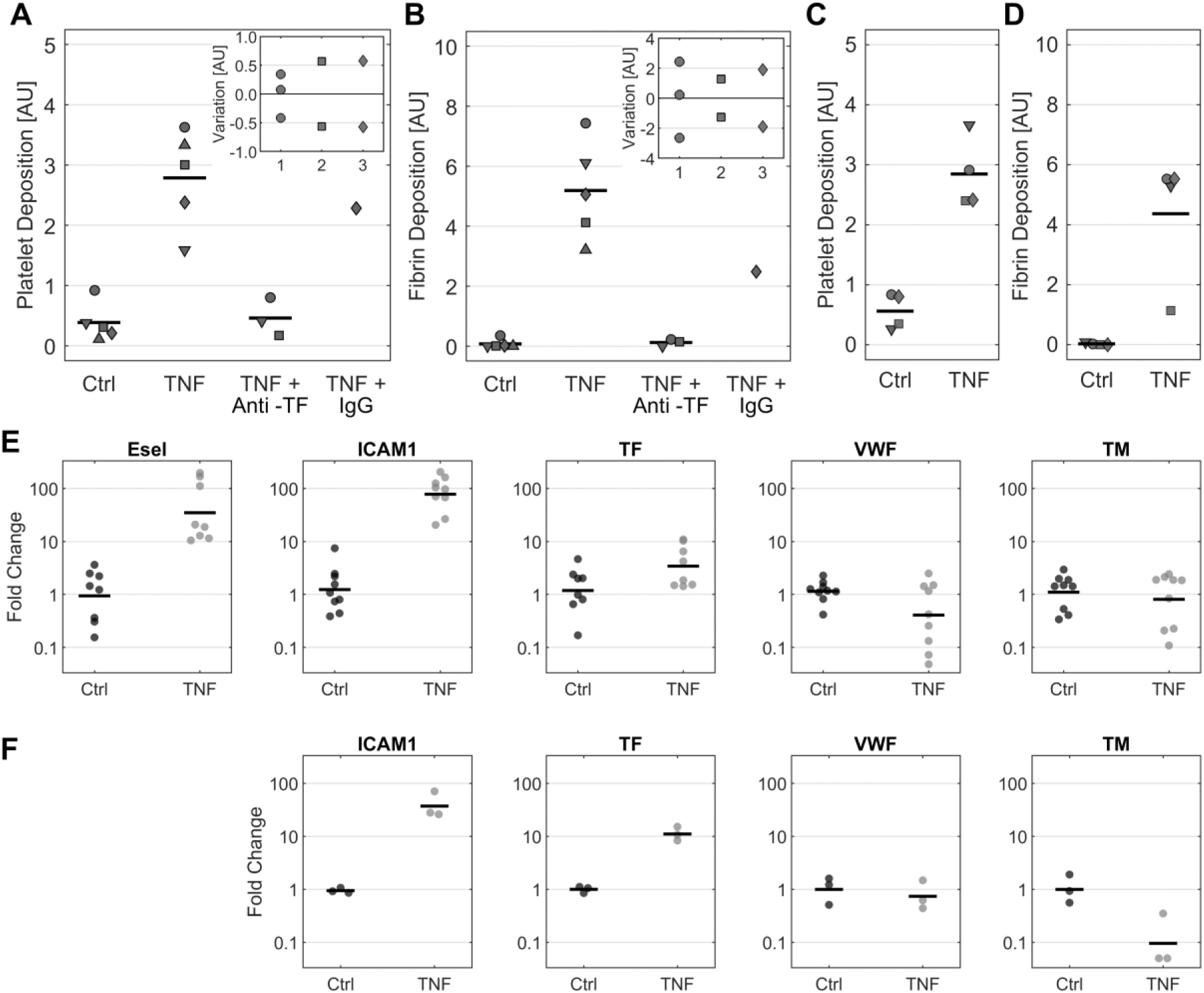
Quantification of thrombus formation and endothelial activation. A-D: Quantification of platelet adhesion and fibrin deposition in HUVEC-lined vessels (A-B) and ECFC-lined vessels (C-D) after 20 min perfusion with healthy whole blood. For both cell types, TNF-α treatment significantly increased platelet adhesion (p<0.01) and fibrin deposition (HUVEC: p<0.001, ECFC: p<0.01). In HUVEC-lined vessels, anti-TF treatment completely abolished both platelet and fibrin deposition, whilst IgG control had no effect. Each symbol represents a single blood donor; when the same blood sample was tested on multiple vessels, the mean value is shown in the main graph and the repeats are shown in the inset panel [Ctrl (1), TNF (2) and TNF + Anti-TF (3)]. E-F: Assessment of endothelial activation by RNA profiling of ICAM1, VWF, TM, TF and E-sel (the latter for HUVEC only). HUVEC vessels (n = 9): TNF-α treatment resulted in a significant upregulation of ICAM1 and E-sel gene expression (p<0.001), a trend towards increase for TF which did not reach statistical significance (p>0.05) and no changes in VWF and TM. ECFC vessels (n = 3): TNF-α treatment induced an upregulation of ICAM1 and TF gene expression (p<0.01); TM was downregulated in all vessels but the sample size was insufficient to conclude a statistical difference (p>0.05); VWF expression was not affected.

TF expression on ECs is upregulated by TNF-α [31]. Thus, an anti-TF blocking antibody (25 µg/ml) was added to the vessel 20 min prior to blood perfusion to evaluate the contribution of TF to thrombus formation in this assay. Figure 2B shows representative images of platelet adhesion and fibrin deposition upon TNF-stimulated HUVEC in the presence of anti-TF or IgG isotype control antibodies. Blocking TF reduced fibrin deposition compared to IgG control or TNF-α alone (p<0.01) (Figure 3B), confirming the involvement of TF in this process. Interestingly, blocking TF also resulted in inhibition of platelet deposition (Figure 3A), suggesting a possible indirect effect from thrombin inhibition, or a contribution of platelet TF.

#### ECFC-lined vessels

Vessels lined with pooled ECFC from healthy donors were used to evaluate the potential for disease modelling. Images show that TNF-α treatment increased platelet adhesion and fibrin deposition compared to untreated controls (representative images in Figure 2C, quantification in Figure 3C-D; p<0.01 for both), comparable to that observed in HUVEC-lined vessels (Figure 3A-B). Platelet deposition was remarkably similar between experiments; however, variability in fibrin deposition was observed, again pointing to vessel variability.

### Validation of endothelial activation by RNA profiling

To verify endothelial activation following TNF-α treatment, gene expression was quantified with qRT-PCR, in HUVEC (n=9, Figure 3E) and ECFC (n=3, Figure 3F). As expected, TNF-α treatment resulted in a significant upregulation of ICAM1 and E-selectin in HUVEC (37-fold and 63-fold on average respectively, p<0.001). Average TF gene expression was upregulated by TNF-α (2.9-fold) but no statistical difference was detected; no differences in VWF and TM mRNA levels were detected (p>0.05). In ECFC, TNF-α stimulation also resulted in upregulation of ICAM1 (39-fold, p<0.01), and TF, the latter with a stronger response compared to HUVEC (11-fold, p<0.01). TNF also caused a 10-fold reduction in TM on average; however, this was not statistically different from zero (p>0.05) likely due to the small sample size. No change in VWF gene expression was detected. These results confirm activation of EC by TNF-α in a 3D vessel structure surrounded by ECM.

### Assessment of endothelial barrier function

Given the variability observed in some repeat samples, we set out to confirm the functional integrity of the monolayers. Barrier function was determined over time by perfusing fluorescent dextran solution through the vessels and estimating monolayer permeability. Confocal fluorescent images taken after 5 min of perfusion (Figure 4A-B) show a reduction in the amount of solute diffusing across the endothelium at Day 9 compared to Day 3. Fluorescence intensity profiles (Figure 4C-D) confirm this trend. At Day 3 (Figure 4A and C), fluorescent dextran was readily transported across the vessel wall, with an increase over the 5 min interval, indicating that the endothelial barrier function has not yet been established. At Day 9 (Figure 4B and D), minimal dextran transport is shown by the low intensity of fluorescent signal in the collagen around the vessel, indicating a tight endothelial barrier.

**Figure 4:**
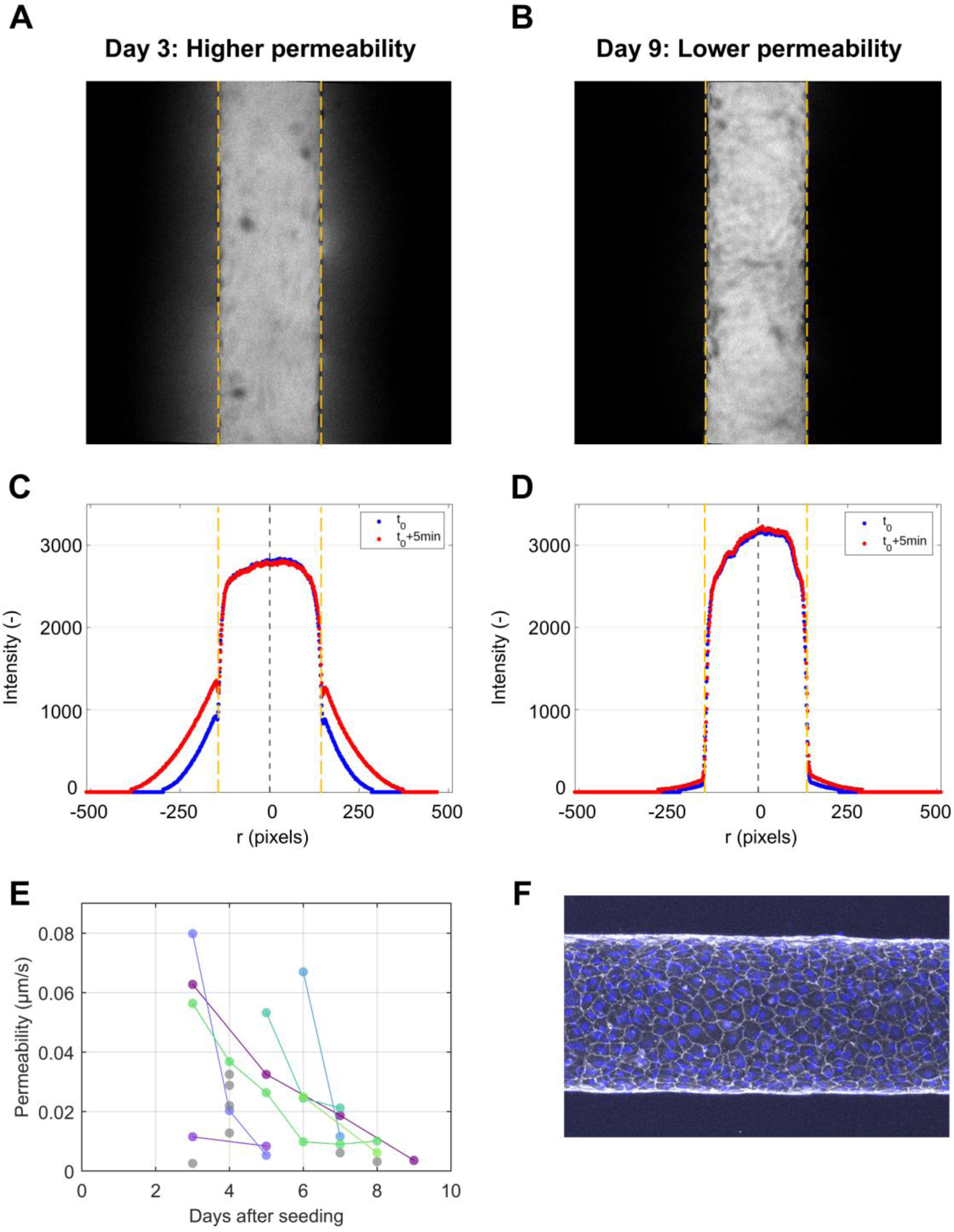
Permeability. Functional evaluation of endothelial barrier tightness. Permeability assay: Dextran diffusion traced after 5 min from dextran homogeneous diffusion in the channel lumen. Confocal images and relative fluorescence profile plotted across the vessel wall of the same microvessel monitored at Day 3 (A and C) and at Day 9 (B and D) of culture. At Day 3, dextran leaked through the monolayer, as sign of an immature endothelial barrier. At Day 9, dextran was completely retained inside the lumen, proving the tightness of the monolayer and a well-established barrier. E: Permeability monitored multiple times across different days in the same vessel (each vessel represented by a different colour) or as individual measurements (grey) in additional vessels is reported; time course data show a decrease in permeability over time for all vessels. Permeability values are comparable to previous reports. F: Confocal imaging: immunostaining against DAPI (blue) and VE-cadherin (white), representing a characteristic confluent microvessel, without visible gaps.

In a subset of vessels, permeability was monitored longitudinally (Figure 4E). A consistent increase in barrier function over time in all vessels was observed; however, variability in the time required by different vessels to achieve a barrier function was observed. The measured permeability values are consistent with previous reports in comparable models (0.001-0.06 µm/s [28], [32]IF staining for junctional VE-Cadherin of a representative vessel at day 5 confirms that a tight barrier corresponds to a confluent endothelial monolayer without visible gaps (Figure 4F, permeability quantified as 0.013 μm/s).

### Co-culture with pericytes

In the microvasculature, the endothelium is surrounded by mural cells, specifically pericytes in capillaries and vascular smooth muscle cells around arterioles and venules. Mural cells wrap around EC and are exposed to blood following vascular injury. Recently pericytes were shown to express TF, thus able to initiate and propagate the extrinsic pathway of the coagulation cascade [33]. The contribution of pericytes to endothelial activation and thrombosis in intact vessels is still largely unknown. We tested whether our new platform was suitable for co-culture of EC with pericytes. Confocal microscopy showed an endothelial monolayer with established continuous junction and pericytes wrapped around the endothelium with thin cellular protrusions around the vessel (Supplementary Figure 1). These images demonstrate the viability of this approach, although the density of pericytes was variable and requires further optimisation.

## DISCUSSION

In this paper, we report the development of an *in vitro* 3D model of thrombosis-on-chip in a controlled microenvironment, using human endothelial cells and whole blood. We demonstrated that the model can be used to study thrombus formation in real time over TNF-α activated endothelium, using blood with fluorescently labelled components and real-time confocal imaging. The model also allows for molecular analysis of endothelial gene expression and incorporation of pericytes and therefore has the potential to incorporate all key components of haemostasis: EC, blood factors, blood cells, ECM, shear stress and perivascular cells. The design supports both biological assays (i.e. immunostaining and real-time qPCR) and functional assays (i.e. permeability and thrombosis assays).

The design of this chip presents several advantages with respect to existing microfluidic models of thrombosis. Contrary to many other devices, the emphasis of our model is on the endothelium: this design is particularly suited to study pathological activation of the endothelium by inflammatory agents and shear stress. Several microfluidic models are fabricated using methods that yield rectangular cross sections [15], [16], [34]. This results in a non-uniform distribution of shear stress on the ECs that could lead to EC activation. To avoid this, we used the needle casting approach, which generates a circular cross-section that provides a uniform shear stress; this method also has the advantage of being easier and cheaper than other microfabrication techniques [18]. Furthermore, the use of inlet and outlet ports that are in-line with the endothelialised vessel are particularly suitable for blood perfusion, as these avoid sharp corners that could potentially induce high local shear-stresses that could activate platelets during perfusion. The model presents an alternative to designs that have incorporated geometries to induce a ‘pathological’ flow profile [35] which result in non-uniform distributions of shear stress. Our system uses the simpler approach of having a single shear stress condition for each vessel that can be controlled via a syringe pump or alternative perfusion device. This provides the flexibility to investigate the different shear stresses associated with venous and arterial thrombosis in a controlled manner.

Endothelial activation by TNF-α was confirmed by the regulation of well-characterised gene targets ICAM1 and E-selectin. Upregulation of TF was observed in ECFC, and the average TF expression was increased in HUVEC, although a statistical difference was not detected. Similarly, the anticoagulant TM was significantly downregulated following TNF-α treatment in ECFCs but not HUVEC; more studies will be required to determine differences between HUVEC and ECFC in response to activating agents. Overall, the response of TNF-α target genes was variable between samples, possibly reflecting a technical issue due to the low quantity of RNA that can be extracted from each vessel. Pooling of multiple vessels could reduce variability. Other endothelial activating agents could be used, including cocktails of pro-inflammatory cytokines for disease modelling of thrombosis.

The model allows manipulation of the coagulation response from both the endothelium and blood components, thus presenting an opportunity to dissect novel pathways and investigate the crosstalk between cellular and circulating regulators. By using anti-TF antibodies, we show that the model is suitable for pathway detection. Our data show that ECFC perform similarly to HUVEC in supporting platelet adhesion and thrombus formation. This is the first step towards developing a personalised model of thrombosis, a key tool to investigate the mechanisms underlying thrombosis of unknown causes. ECFCs are a unique tool to create an autologous model of thrombosis, by pairing ECFCs with blood isolated from the same patient for pathway investigation and drug testing.

Another key feature of our device is the direct interaction between EC and the surrounding ECM, thus allowing the cells to sense the substrate stiffness and crucially engage in signals from the ECM. By changing collagen concentration or ECM substrate, future studies could investigate the importance of stiffness in the response of the endothelium to activation and the consequent thrombus formation.

Finally, our vessel-on-a-chip platform can be used to investigate the crosstalk between EC and pericytes, embedded in the same ECM and thus able to engage in direct cell-cell interactions without separation by a barrier; this would not be possible using membrane-based co-culture or microfluidic models.

There are limitations to this study. While the needle-casting fabrication method provides the advantages noted above, it is practically limited to vessel scales on the order of 100µm or larger and straight cylindrical geometries. Nonetheless, given the need for parallelisation to achieve sufficient statistical power for an assay, the addition of complexity could hinder the practical use of the assay. We observed variability between the vessels, which could be due to several elements. By testing the same blood sample on multiple vessels in the same experiment, we concluded that some variability could be attributed to the vessels. A low but detectable level of platelet adhesion was detected on non-activated vessels. This was not a major issue for model evaluation, as TNF-α stimulation resulted in >3-fold increase in platelet adhesion compared to baseline. However, the presence of background platelet adhesion could become a limitation if a less powerful inflammatory stimulation is used, or to detect spontaneous platelet adhesion. Although platelets activation state could be characterised prior to this assay to provide a baseline assessment of donor variability, there is a practical time limitation as blood samples should be used within hours of collection [36]. Since close to zero fibrin deposition could be detected in unstimulated vessels, this suggests that the background level of platelet adhesion in unstimulated vessels was not thrombin-dependent, but rather that the platelets were already activated before entering the vessel, as a consequence of sample manipulation. Further optimisation will be required to reduce this background.

The value of using live-imaging to study thrombus formation has been extensively demonstrated *in vivo* [37]. The flexibility in labelling different blood components could allow to investigate the contribution of other cells, i.e. leukocytes, to thrombus formation. Time-lapse imaging enabled the dynamic response of platelet adhesion and fibrin deposition to be captured; optimisation experiments showed that 20 minutes imaging time was sufficient to capture differences in coagulation between the two treatment groups (control and TNF-α). Using z-stacks was important to capture the surface area of the vessel; however, the current method for data analysis relies on the platelet channel for defining the vessel boundaries. This might result in an underestimation of the vessel boundaries, especially in the control vessels with lower platelet coverage. Live labelling of ECs could more accurately define vessel boundaries; this would provide higher accuracy for determining the start point of the z-stack, thereby reducing the number of images required per z-stack, which would reduce the imaging time. However, reagents used for live cell staining are often toxic to cells. More reliable methods for fluorescent labelling, including lentivirus, can reduce cell viability in long-term culture and would be practically inconvenient to implement with patients’ ECFCs.

In conclusion, we have developed a vessel-on-a-chip microfluidic model of thrombosis that provides flexibility to manipulate multiple variables relevant to thrombus formation within a 3D vessel structure. This model provides a platform to dissect the mechanisms of thrombus formation, investigate the effect of coagulation inhibitors, predict thrombotic side effects of drugs and study the pathological mechanisms underlying thrombus formation in patients with thrombosis of unknown causes, using patients’ own EC and blood in an autologous, precision medicine approach.

## AUTHORSHIP DETAILS

Conceptualisation: JJE, AMR, JvBS; Experiments: JJE, MD, TM; CP; Data analysis: JJE, TM, JvBS. Experimental and scientific advice: JA; CP; CPer. Funding AMR, JvBS. Writing the manuscript: AMR, TM, JJE, JvBS. Critical revision: all authors.

## ACKNOWLEDGEMENTS

This work was supported by Imperial College NHLI-British Heart Foundation (BHF) PhD studentship (JJE); BHF programme grant RG/17/4/32662 (MD); the National Institute for Health Research (NIHR) Imperial Biomedical Research Centre Cardiovascular Theme (AMR; CPer); Imperial BHF Research Excellence Award (4) – RE/24/130023 (AR); PNRR PNC-E3-2022-23683266 PNC-HLS-DA, INNOVA (TM, AR); Royal Academy of Engineering Research Fellowship RF201617\16\18 (JvBS); Versus Arthritis Career Development Fellowship 21223 and the Imperial College–Wellcome Trust Institutional Strategic Support Fund (CPer). We also thank the BHF Center of Excellence for support (AMR). We thank Dr. Daniela Pirri, Dr. Tom McKinnon and Dr. Graeme Birdsey (Imperial College London) for technical support.

## Supplementary data

**Suppl Figure 1:**
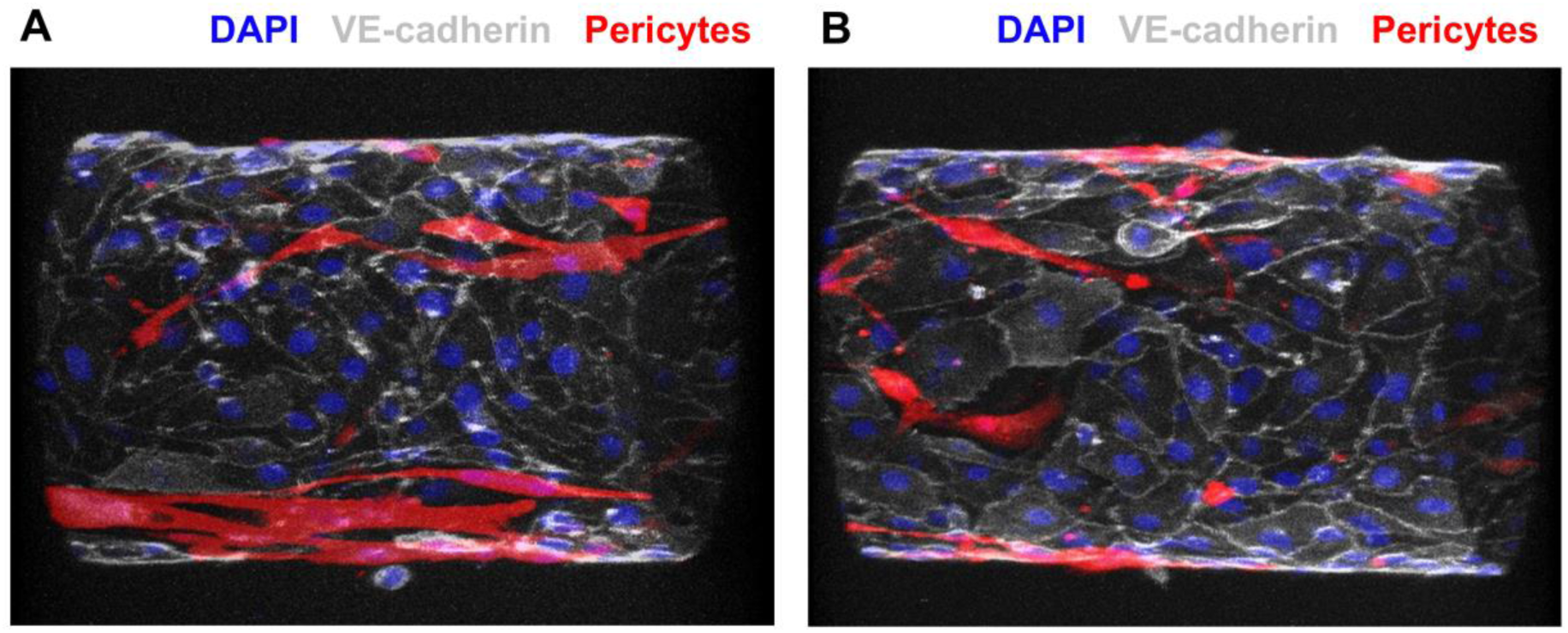
co-culture with pericytes. Confocal imaging of vessels co-cultured with HUVEC and HBMVP immunostained against DAPI (blue) and VE-cadherin (white) and with RFP-labelled pericytes (red). Images show the formation of an endothelial monolayer in the presence of pericytes, demonstrating the viability of the method for co-culture studies. Variability in pericyte coverage was observed between different vessels and requires further optimisation. Scale bars = 100 µm

## REFERENCES

[1] H. H. Versteeg, J. W. M. Heemskerk, M. Levi, and P. H. Reitsma, “New Fundamentals in hemostasis,” Jan. 01, 2013, American Physiological Society Bethesda, MD. doi: 10.1152/physrev.00016.2011.

[2] N. Mackman, R. E. Tilley, and N. S. Key, “Role of the extrinsic pathway of blood coagulation in hemostasis and thrombosis,” Arterioscler Thromb Vasc Biol, vol. 27, no. 8, pp. 1687–1693, Aug. 2007, doi: 10.1161/ATVBAHA.107.141911.

[3] T. G. Mastenbroek, J. P. van Geffen, J. W. M. Heemskerk, and J. M. E. M. Cosemans, “Acute and persistent platelet and coagulant activities in atherothrombosis,” J Thromb Haemost, vol. 13 Suppl 1, no. S1, pp. S272–S280, Jun. 2015, doi: 10.1111/JTH.12972.

[4] B. Furie and B. C. Furie, “Mechanisms of Thrombus Formation,” N Engl J Med, 2008.

[5] E. M. Van Cott, B. Khor, and J. L. Zehnder, “Factor V Leiden,” Am J Hematol, vol. 91, no. 1, pp. 46–49, Jan. 2016, doi: 10.1002/AJH.24222.

[6] C. Zuchi, I. Tritto, E. Carluccio, C. Mattei, G. Cattadori, and G. Ambrosio, “Role of endothelial dysfunction in heart failure,” 2020. doi: 10.1007/s10741-019-09881-3.

[7] P. Poredos and M. K. Jezovnik, “Endothelial Dysfunction and Venous Thrombosis,” Sep. 27, 2018, SAGE PublicationsSage CA: Los Angeles, CA. doi: 10.1177/0003319717732238.

[8] M. A. Gimbrone and G. García-Cardeña, “Endothelial Cell Dysfunction and the Pathobiology of Atherosclerosis,” Circ Res, vol. 118, no. 4, pp. 620–636, 2016, doi: 10.1161/CIRCRESAHA.115.306301.

[9] P. Libby, “Inflammation in atherosclerosis,” Arterioscler Thromb Vasc Biol, vol. 32, no. 9, pp. 2045–2051, Sep. 2012, doi: 10.1161/ATVBAHA.108.179705.

[10] B. L. Doss et al., “Cell response to substrate rigidity is regulated by active and passive cytoskeletal stress,” Proc Natl Acad Sci U S A, vol. 117, no. 23, pp. 12817–12825, 2020, doi: 10.1073/pnas.1917555117.

[11] R. R. Rao, A. W. Peterson, J. Ceccarelli, A. J. Putnam, and J. P. Stegemann, “Matrix composition regulates three-dimensional network formation by endothelial cells and mesenchymal stem cells in collagen/fibrin materials,” Angiogenesis, vol. 15, no. 2, pp. 253–264, 2012, doi: 10.1007/s10456-012-9257-1.

[12] C.-M. Lo, H.-B. Wang, M. Dembo, and Y.-L. Wang, “Cell Movement Is Guided by the Rigidity of the Substrate,” Biophys J, 2000, doi: 10.1016/S0006-3495(00)76279-5.

[13] M. Sweeney and G. Foldes, “It Takes Two: Endothelial-Perivascular Cell Cross-Talk in Vascular Development and Disease,” Front Cardiovasc Med, vol. 5, no. October, pp. 1–14, 2018, doi: 10.3389/fcvm.2018.00154.

[14] A. Geevarghese and I. M. Herman, “Pericyte-Endothelial Cross-Talk: Implications and Opportunities for Advanced Cellular Therapies,” Transl Res, vol. 163, no. 4, p. 296, 2014, doi: 10.1016/J.TRSL.2014.01.011.

[15] T. Mathur et al., “Organ-on-chips made of blood: Endothelial progenitor cells from blood reconstitute vascular thromboinflammation in vessel-chips,” Lab Chip, vol. 19, no. 15, pp. 2500–2511, 2019, doi: 10.1039/c9lc00469f.

[16] X. D. Manz et al., “In vitro microfluidic disease model to study whole blood-endothelial interactions and blood clot dynamics in real-time,” Journal of Visualized Experiments, vol. 2020, no. 159, pp. 1–10, 2020, doi: 10.3791/61068.

[17] S. Ayyoub, R. Orriols, E. Oliver, and O. T. Ceide, “Thrombosis Models: An Overview of Common In Vivo and In Vitro Models of Thrombosis,” Feb. 01, 2023, *MDPI*. doi: 10.3390/ijms24032569.

[18] A. Shakeri et al., “Engineering Organ-on-a-Chip Systems for Vascular Diseases,” Dec. 01, 2023, Lippincott Williams and Wilkins. doi: 10.1161/ATVBAHA.123.318233.

[19] J. W. Weisel and R. I. Litvinov, “Red blood cells: the forgotten player in hemostasis and thrombosis,” Feb. 01, 2019, Blackwell Publishing Ltd. doi: 10.1111/jth.14360.

[20] Y. Qiu et al., “Clinically relevant clot resolution via a thromboinflammation-on-a-chip,” Nature, 2025, doi: 10.1038/s41586-025-08804-7.

[21] Y. C. Zhao et al., “Novel Movable Typing for Personalized Vein-Chips in Large Scale: Recapitulate Patient-Specific Virchow’s Triad and its Contribution to Cerebral Venous Sinus Thrombosis,” Adv Funct Mater, vol. 33, no. 23, Jun. 2023, doi: 10.1002/adfm.202214179.

[22] C. Cuartas-Vélez, H. H. T. Middelkamp, A. D. van der Meer, A. van den Berg, and N. Bosschaart, “Tracking the dynamics of thrombus formation in a blood vessel-on-chip with visible-light optical coherence tomography,” Biomed Opt Express, vol. 14, no. 11, p. 5642, Nov. 2023, doi: 10.1364/boe.500434.

[23] R. Barrile et al., “Organ-on-Chip Recapitulates Thrombosis Induced by an anti-CD154 Monoclonal Antibody: Translational Potential of Advanced Microengineered Systems,” Clin Pharmacol Ther, vol. 104, no. 6, pp. 1240–1248, Dec. 2018, doi: 10.1002/cpt.1054.

[24] Y. Sakurai et al., “A microengineered vascularized bleeding model that integrates the principal components of hemostasis,” Nat Commun, vol. 9, no. 1, Dec. 2018, doi: 10.1038/s41467-018-02990-x.

[25] P. F. Costa et al., “Mimicking arterial thrombosis in a 3D-printed microfluidic: In vitro vascular model based on computed tomography angiography data,” Lab Chip, vol. 17, no. 16, pp. 2785–2792, Aug. 2017, doi: 10.1039/c7lc00202e.

[26] K. E. Paschalaki and A. M. Randi, “Recent advances in endothelial colony forming cells toward their use in clinical translation,” Front Med (Lausanne), vol. 5, no. OCT, pp. 1–12, 2018, doi: 10.3389/fmed.2018.00295.

[27] S. Della Bella et al., “Pathologic up-regulation of TNFSF15-TNFRSF25 axis sustains endothelial dysfunction in unprovoked venous thromboembolism,” Cardiovasc Res, vol. 116, no. 3, pp. 698–707, 2020, doi: 10.1093/cvr/cvz131.

[28] K. M. Chrobak, D. R. Potter, and J. Tien, “Formation of perfused, functional microvascular tubes in vitro,” Microvasc Res, vol. 71, no. 3, pp. 185–196, May 2006, doi: 10.1016/j.mvr.2006.02.005.

[29] R. D. Starke et al., “Cellular and molecular basis of von Willebrand disease: studies on blood outgrowth endothelial cells Key Points • BOECs from VWD patients provide novel insight into the cellular mechanisms of the disease,” Blood, vol. 121, no. 14, pp. 2773–2784, 2013, doi: 10.1182/blood-2012-06.

[30] T. Mathur et al., “Organ-on-chips made of blood: Endothelial progenitor cells from blood reconstitute vascular thromboinflammation in vessel-chips,” Lab Chip, vol. 19, no. 15, pp. 2500–2511, 2019, doi: 10.1039/c9lc00469f.

[31] D. Kirchhofer, T. B. Tschopp, P. Hadvary, and H. R. Baumgartner, “Endothelial cells stimulated with tumor necrosis factor-α express varying amounts of tissue factor resulting in inhomogenous fibrin deposition in a native blood flow system. Effects of thrombin inhibitors,” Journal of Clinical Investigation, vol. 93, no. 5, pp. 2073–2083, 1994, doi: 10.1172/JCI117202.

[32] G. S. Offeddu et al., “An on-chip model of protein paracellular and transcellular permeability in the microcirculation,” Biomaterials, vol. 212, no. April, pp. 115–125, 2019, doi: 10.1016/j.biomaterials.2019.05.022.

[33] B. A. Bouchard, M. A. Shatos, and P. B. Tracy, “Human Brain Pericytes Differentially Regulate Expression of Procoagulant Enzyme Complexes Comprising the Extrinsic Pathway of Blood Coagulation,” Arterioscler Thromb Vasc Biol, vol. 17, no. 1, pp. 1–9, 1997, doi: 10.1161/01.ATV.17.1.1.

[34] A. Jain et al., “Assessment of whole blood thrombosis in a microfluidic device lined by fixed human endothelium,” Biomed Microdevices, vol. 18, no. 4, pp. 1–7, 2016, doi: 10.1007/s10544-016-0095-6.

[35] N. V. Menon et al., “Recapitulating atherogenic flow disturbances and vascular inflammation in a perfusable 3D stenosis model,” Biofabrication, vol. 12, no. 4, Oct. 2020, doi: 10.1088/1758-5090/ABA501.

[36] O. K. Baskurt et al., “New guidelines for hemorheological laboratory techniques,” Clin Hemorheol Microcirc, vol. 42, no. 2, pp. 75–97, 2009, doi: 10.3233/CH-2009-1202.

[37] B. Furie and B. Furie, “In vivo thrombus formation,” J Thromb Haemost, vol. 5, pp. 12–17, 2007.

