## Appendices for "A Microfluidic Blood Vessel-On-Chip Model of Thrombosis"

### A1 Design of the perfusion unit

#### Design criteria

There are two design criteria that influence the design of the perfusion unit designed for the culture and maturation of the endothelium of the vessel:

1. Media turnover rate should be sufficiently high to ensure adequate nutrient delivery to cells.
2. Shear stress applied to the endothelial cells should be as consistent as possible. For this study, we aim to maintain a low shear stress that will not modify cell behaviour.

Media turnover rate: for a cylindrical vessel of radius $R$ and length $L$, the turnover rate $f_V$ in vessel volumes per minute is given by

$$f_V = \frac{Q}{\pi R^2 L} \quad \text{(A1-1)}$$

Shear stress: for a cylindrical vessel the shear stress $\tau$ is directly related to the flow rate $Q$ according to

$$\tau = \frac{4\mu}{\pi R^3} Q \quad \text{(A1-2)}$$

where $\mu$ is the viscosity of the cell culture media. We wish to avoid $\tau$ changing significantly during culture, and hence $Q$ should vary as little as possible.

#### Flow versus pressure control

The dependency on flow rate of both shear stress and media turnover implies that a flow-based control system would be optimal. Peristaltic pumps can provide flow control, but generate oscillatory flow rates that have to be damped with additional fluidic components for each channel. Syringe pumps deliver more stable flow, although oscillations are still present and there is a trade-off between the syringe size and the stability of the flow. Both systems require individual pump channels and additional tubing per vessel, making parallelised unwieldy and expensive.

Hydrostatic pressure offers a practical alternative that is a lot simpler to implement, is low cost and parallelisable. For a long-straight vessel, the Hagen-Poiseuille equation relates $Q$ to the pressure drop driving the flow, $\Delta p$, according to

$$Q = \frac{\pi R^4}{8\mu}\frac{\Delta p}{L} = C_H \Delta p \quad \text{(A1-3)}$$

The term $C_H$ represents the hydrodynamic conductance of the vessel. Assuming the vessel dimensions and hence $C_H$ are fixed by fabrication considerations, $Q$ is directly proportional to $\Delta p$. Hence we aim to minimise changes in $\Delta p$ during the culture phase.

#### Reduction in hydrostatic pressure drop over time

The hydrostatic pressure driving fluid movement through a single vessel is given by

$$\Delta p = \rho g (h_i - h_o) \quad \text{(A1-4)}$$

where $\rho$ is the density of the cell culture media, $g$ is gravitational acceleration, and $h_i$ and $h_o$ are the heights of the fluid in the inlet and outlet reservoirs respectively (see Figure 1E).

As media flows from the inlet reservoir at flow rate $Q$, the volume of fluid in the reservoir decreases, and correspondingly so does the height of the fluid in the reservoir,

$$Q(t) = \frac{dV_i}{dt} = -A\frac{dh_i}{dt} \tag{A1-5}$$

where $A$ is the area of the fluid surface in the inlet reservoir. Due to conservation of mass, the volume of the outlet reservoir increases by the same amount, and assuming the same cross sectional area $A$:

$$Q(t) = \frac{dV_o}{dt} = A\frac{dh_o}{dt} \tag{A1-6}$$

As the heights change, so does the pressure driving the flow. By differentiating Equation A1-4, we see that

$$\frac{d(\Delta p(t))}{dt} = \rho g\left(\frac{dh_i}{dt} - \frac{dh_o}{dt}\right) = -\frac{2\rho g Q(t)}{A} \tag{A1-7}$$

Or in terms of flow rate via Equation A3

$$\frac{dQ}{dt} = -\frac{2\rho g}{AC}Q(t) \tag{A1-8}$$

These changes in $Q$ can be countered by topping up the inlet reservoir and emptying out the outlet reservoir, but for the practical use of the system, this should be necessary at most once per day.

**Minimising changes in flow rate**

We aim to minimise $dQ/dt$ in order to minimise changes to $Q$. Decreasing the media density $\rho$ is not practical. It would be possible to increase the media viscosity $\mu$, which would decrease conductivity, $C_H$, and hence $dQ/dt$. However, this would require validation for each cell type that it is not affecting cell function. Therefore, maximising $A$ is the only practical approach to reducing $dQ/dt$, although there are trade-offs with practicality as the culture unit needs to be transportable from the incubator to the hood for permeability measurements and visual checks.

To further decrease $dQ/dt$, we added an additional design feature to the culture unit. The outlet reservoirs were designed with an overflow unit, such that $dh_o/dt$ was constant. This halves the rate of change of $Q$ to

$$\frac{dQ}{dt} = -\frac{\rho g}{AC}Q(t) \tag{A1-9}$$

For the current design, with a media turnover rate of once per day, we achieved $f_V$>1 channel volume per minute and $\tau < 0.3\ dyne/cm^2$.

### A2 Measurement of Permeability
#### Theory

Confocal images were captured with a constant focal depth $f$ over time following infusion of fluorescent dye into the vessel. Permeability analysis was performed using MATLAB, based on the assumption of radial diffusion from a cylinder (Figure A1).

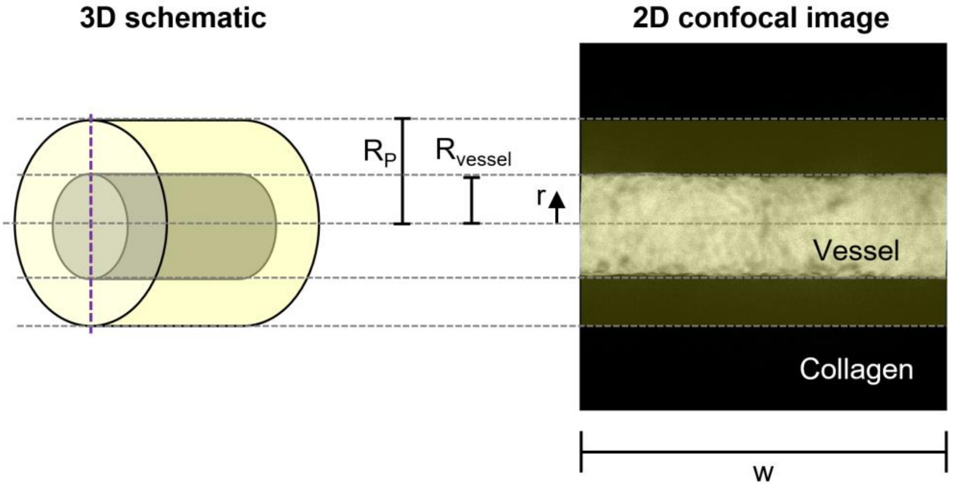


**Figure A1**: schematic of the cylinder and corresponding image, with dimensions indicated.

The permeability, $P_s$, of a specific solute across an endothelial barrier is defined according to

$$P_s = \frac{J_s}{S\Delta C} \quad \text{(A2-1)}$$

where $J_s$ is the solute flux across the vessel wall, $S$ is the surface area of the vessel and $\Delta C$ is the concentration gradient driving the diffusion of the solute.

$$S = 2\pi R_{vessel} w \quad \text{(A2-2)}$$

$$\Delta C(t) = C_{vessel} - C_{ECM}(t) \approx \overline{C_{vessel}} \quad \text{(A2-3)}$$

Where $C_{vessel}$ is the concentration in the vessel (assumed to be constant) and $C_{ECM}(t)$ is the concentration in the ECM outside the vessel (assumed to be uniform). Further assuming that $C_{ECM}(t) \ll C_{vessel}$, we can neglect the second term, simplifying the analysis. Further, we assume that our region of interest is sufficiently small that we can ignore spatial distribution of $C_{ECM}$, and hence use $\overline{C_{ECM}}(t)$.

$J_s$ is the number of moles transported in a unit time out of the vessel, and is given by

$$J_s(t) = \frac{dn_{ECM}(t)}{dt} = V_{ECM}\frac{d\overline{C_{ECM}}(t)}{dt} \quad \text{(A2-4)}$$

The volume is given by

$$V_{ECM} = \pi(R_P^2 - R_{vessel}^2)w \quad \text{(A2-5)}$$

This leaves $dC_{ECM}(t)/dt$ to calculate, which can be done via fluorescent confocal imaging, under the assumption that the fluorescent intensity is linearly proportional to the number of moles of fluorescent Dextran in the imaging volume, such that

$$C = \alpha I \quad \text{(A2-6)}$$

where $\alpha$ is a constant that can be acquired via calibration of fluorescent intensity against a known concentration (but cancels out).

Together, Equations A2-1 to 6 yield

$$P_s = \frac{R_P^2 - R_{vessel}^2}{2R_{vessel}} \frac{1}{(\overline{I_{vessel}})} \frac{d\overline{I_{ECM}}(t)}{dt} \quad \text{(A2-7)}$$

**Implementation**

Averaging along the axial length of the vessel $(w)$ yields an average profile of image intensity (Figure 4C), from which $R_{vessel}$ can be extracted. $R_P$ was selected to not include any zero-intensity pixels in the ECM and should be small enough for the assumption of uniformity to not introduce significant error. By inspection of all data sets,

$$R_P = 4/3R_{vessel} \quad \text{(A2-8)}$$

was selected for permeability analysis of all vessels.

To ensure the assumption of constant $C_{vessel}$ was achieved, 'time zero' was selected as the earliest time that the value of $I_{vessel}$ changes by less than 5% over a five minute window, after increasing from zero at the beginning of the perfusion. $\overline{I_{vessel}}$ was the average intensity in the vessel over those 5 minutes, and $\frac{d\overline{I_{ECM}}(t)}{dt}$ was calculated by fitting a straight line to the average intensity in the ECM region over time.

### A3 Measurement of Platelet Adhesion and Fibrin Deposition

#### Theory

For quantification of platelet adhesion and fibrin deposition,14 images were captured for each vessel and assembled as z-stacks (Figure A2).

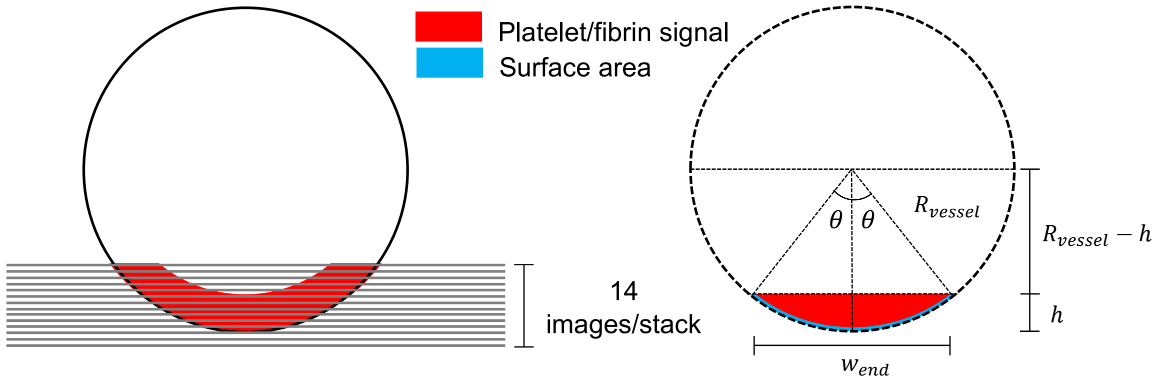


**Figure A2**: schematic of the cross section of the vessel and platelet/fibrin signal for normalisation.

As the imaged area can vary between vessels, we normalised the total signal by the vessel area captured:

$$A_{surface} = 2R_{vessel}l\theta \tag{A3-1}$$

where $l$ is the length of the image in the flow direction. The value of $\theta$ can be calculated according to

$$\theta = \sin^{-1}\frac{w_{end}}{2R_{vessel}} \tag{A3-2}$$

The values of $w_{end}$ and $h$ can be directly calculated from the images, but $R_{vessel}$ cannot. However, Pythagoras' theorem yields

$$R_{vessel} = \frac{w_{end}^2}{8h} + \frac{h}{2} \tag{A3-3}$$

#### Implementation

Image stacks were pre-processed in Fiji (ImageJ) by applying a median despeckle filter, and then analysed in MATLAB (Mathworks, MA, USA). Platelet- or fibrin-occupied area was defined as the area within the vessel boundaries with pixel intensity, $I > 0$ (as the background intensity was 0 for both channels). $w_{end}$ was calculated based on the width of the non-zero signal within the uppermost image of the stack, and $h$ was calculated based on the number of slices between the uppermost image and the first image that contained a signal (which represents the bottom of the vessel). Equations A3-1 to 3 were used to calculate $A_{surface}$ and the normalised intensity for the image was calculated as the sum of the intensity across the stack, divided by $A_{surface}$. This is presented in arbitrary units (AU) as Platelet Deposition [AU] or Fibrin Deposition [AU] in Figure 3.

### A4 Statistical Analysis

The data sets in this study comprise a mix of variables and readouts, but a full balanced experimental design was not practical, due to the technical complexity of the experiments. For simplicity and interpretability, for each analysis of interest, we use pair-wise analyses and correct for multiple comparisons using the Bonferonni-Holm approach. This reduces the likelihood of a false positive.

Due to the small sample sizes, particularly for the ECFC (n=3 or n=4), the non-parametric Mann-Whitney U-test was not an option, so we instead must rely on unpaired t-tests. Accordingly, we test each sample for the null hypothesis of each sample analysed having arisen from a lognormal or normal distribution. In each data set, only a few samples had p-values that implied the data may not be sampled from a normal distribution (listed below for completeness). In the interest of practicality, we considered this an acceptable uncertainty for use of the t-test.

**Statistical analysis for Platelet Adhesion and Fibrin Deposition**

Blood samples from 5 or 4 individuals were perfused through HUVEC- or ECFC-lined vessel respectively. For each donor, 1-3 vessels were used and outcomes for a given donor were averaged across vessels, although all data points are visualised. For both cell types, untreated (control) and TNF-treated blood was evaluated for all vessels (n=5 and n=4 for HUVEC and ECFC respectively), and we evaluated the effect of TNF on the platelet and fibrin levels. Additionally, n=3 HUVEC-lined vessels were perfused with a combination TNF and Anti-TF to evaluate whether the Anti-TF blocked the effect of the TNF.

The Shapiro-Wilk test for the HUVEC Control and ECFC TF fibrinogen data yielded p-values that implied the data were not sampled from a normal distribution ($0.001<p<0.01$), but the other 8 samples yielded $p>0.05$.

Four t-tests were therefore carried to evaluate the effect of TF on platelets and fibrinogen on for HUVEC and ECFC, with a further two tests to evaluate the effect of Anti-TF. The Bonferonni-Holm correction was applied to these 6 tests.

**Statistical analysis for qPCR**

The data sets comprise the results from 5 or 4 genes on control vs TNF-treated vessels. For HUVEC we had n=8 or 9 independent samples and for ECFC, n=3.

As the data are based on powers of 2, we tested the null hypothesis of the samples coming from lognormal distribution using the Shapiro-Wilk test. For HUVECs, two values were in the range $0.01<p<0.05$ and for ECFC a single case had 2 out of 3 data points were identical, so a p-value could not be computed. The remaining were >0.05.

Nine t-tests were carried out on the log-transform of the PCR the effect of TNF for each of the genes and cell types, and the Bonferonni-Holm correction was applied to these 9 tests.

As the data are log-normally distributed, fold changes in gene expression are calculated using the log-transformed values.
